# Fast and accurate protein function prediction from sequence through pretrained language model and homology-based label diffusion

**DOI:** 10.1101/2022.12.05.519119

**Authors:** Qianmu Yuan, Junjie Xie, Jiancong Xie, Huiying Zhao, Yuedong Yang

**Author notes:** Corresponding author: Yuedong Yang, School of Computer Science and Engineering, Sun Yat-sen University, Guangzhou 510000, China, and Key Laboratory of Machine Intelligence and Advanced Computing of MOE, Sun Yat-sen University, Guangzhou 510000, China.; Huiying Zhao, Sun Yat-sen Memorial Hospital, Sun Yat-sen University, Guangzhou 510000, China. Author biography. Qianmu Yuan School of Computer Science and Engineering at Sun Yat-sen University. His research interests lie in deep learning, graph neural network, protein structure prediction, and protein function prediction. Junjie Xie School of Computer Science and Engineering at Sun Yat-sen University. His research interests include deep learning, graph neural network, and molecule generation. Jiancong Xie School of Computer Science and Engineering at Sun Yat-sen University. His research interests include deep learning, graph neural network, and knowledge graph. Huiying Zhao Sun Yat-sen Memorial Hospital at Sun Yat-sen University. Her research interests include pathogenic gene analysis, protein function, and RNA function prediction. Yuedong Yang School of Computer Science and Engineering at Sun Yat-sen University. Currently he focuses on integrating HPC and AI algorithms for biomedical research.

## Abstract

Protein function prediction is an essential task in bioinformatics which benefits disease mechanism elucidation and drug target discovery. Due to the explosive growth of proteins in sequence databases and the diversity of their functions, it remains challenging to fast and accurately predict protein functions from sequences alone. Although many methods have integrated protein structures, biological networks or literature information to improve performance, these extra features are often unavailable for most proteins. Here, we propose SPROF-GO, a Sequence-based alignment-free PROtein Function predictor which leverages a pretrained language model to efficiently extract informative sequence embeddings and employs self-attention pooling to focus on important residues. The prediction is further advanced by exploiting the homology information and accounting for the overlapping communities of proteins with related functions through the label diffusion algorithm. SPROF-GO was shown to surpass state-of-the-art sequence-based and even network-based approaches by more than 14.5%, 27.3% and 10.1% in AUPR on the three sub-ontology test sets, respectively. Our method was also demonstrated to generalize well on non-homologous proteins and unseen species. Finally, visualization based on the attention mechanism indicated that SPROF-GO is able to capture sequence domains useful for function prediction.

**Key points:**

- SPROF-GO is a sequence-based protein function predictor which leverages a pretrained language model to efficiently extract informative sequence embeddings, thus bypassing expensive database searches.
- SPROF-GO employs self-attention pooling to capture sequence domains useful for function prediction and provide interpretability.
- SPROF-GO applies hierarchical learning strategy to produce consistent predictions and label diffusion to exploit the homology information.
- SPROF-GO is accurate and robust, with better performance than state-of-the-art sequence-based and even network-based approaches, and great generalization ability on non-homologous proteins and unseen species

## 1. Introduction

Proteins play crucial roles within living organisms, including signal transduction, catalysis of metabolic reaction, and maintenance of cellular structure. Identification of protein functions benefits disease mechanism elucidation and drug target discovery [1]. Since traditional biochemical experiments to determine protein functions are usually expensive, time-consuming, and of low throughput [2], fewer than 0.1% of the available protein sequences are currently annotated with reliable information [3], and the gap between unannotated and annotated sequences is expanding at an unparalleled rate [4]. Therefore, it is imperative to develop efficient and effective computational methods for protein function prediction [5].

The functions of proteins are standardized by Gene Ontology (GO) [6], which covers three biological domains: molecular function (MF), biological process (BP) and cellular component (CC), with over 43,000 classes/terms (November 2022). Since a protein is usually associated with multiple GO terms, protein function prediction can be regarded as a large-scale, multi-class, multi-label problem. Moreover, GO is a directed acyclic graph (DAG), in which if a protein is annotated with a GO term, all its ancestor terms up to the root of the ontology should also be annotated. Therefore, protein function predictors should take the hierarchical structure of GO into account and yield “consistent” outputs: the predicted probability of a GO term must be equal to or greater than all of its child terms [7]. To facilitate this challenging task, the critical assessment of functional annotation (CAFA) competition has been held four times [5, 8, 9] using a time-delayed evaluation process. Specifically, given the target proteins, participants were required to submit the predictions before T_0_. After a few months (T_1_), the organizers collected proteins with new experimental annotations as the final test set, consisting of no-knowledge and limited-knowledge proteins. Both types of proteins received their first experimental annotations in the target GO domain between T_0_ and T_1_. However, no-knowledge proteins did not have any experimental annotations before T_0_, while limited-knowledge proteins did in domains other than the target domain. Here we focus on the function prediction for no-knowledge proteins as the vast majority of proteins have no experimental annotations.

Current protein function predictors can be roughly grouped into four categories according to their used information: sequence-based, structure-based, biological network-based, and biomedical literature-based methods. Most sequence-based methods employ sequence similarity, search for sequence domains, or adopt deep learning to capture discriminative features to infer functions. Specifically, a basic way is to transfer annotations directly from homologous sequences with known functions, like Blast2GO [10], since similar sequences tend to have similar functions [5]. Another approach is to search for functional sequence domains or families. For example, GOLabeler [11] utilizes learning to rank algorithm to integrate sequence homology, protein domains and families derived from sequences by BLAST [12] and InterProScan [13]. With the development of deep learning technology, discriminative embeddings can also be automatically extracted from preliminary sequences through designing complex neural networks, including convolutional neural networks in DeepGOPlus [14] and transformer in TALE [15]. However, current sequence-based methods suffer from either low predicted accuracy or the high computational cost (owing to the usage of multi-sequence alignment). On the other hand, recent structure-based methods apply native or predicted protein structures as input, usually followed by graph neural networks (GNN) to learn the local tertiary patterns for function prediction, as in DeepFRI [16] and GAT-GO [17]. Network-based methods exploit the rationale that proteins connected in biological networks (e.g. protein-protein interaction (PPI) or metabolic network) are likely to share the same functions [18]. For example, NetGO [19] integrates multiple protein networks in STRING [20] and transfers annotations from nearest neighbors in the aggregated network. DeepGO [21] adopts knowledge graph embedding algorithm to learn protein features from PPI networks. S2F [22] transfers PPI networks from model organisms to newly sequenced ones, in which label diffusion is employed to propagate initial predictions from several sequence-based component predictors. DeepGraphGO [23] makes the most of both protein sequence domain and high-order protein network information via multispecies GNN strategy. Literature-based methods like DeepText2GO [24] attempt to extract explicit descriptions of protein functions or properties from biomedical texts. NetGO 2.0 [25] incorporates literature and latent sequence information into NetGO to further improve performance. Although CAFA challenge has shown that integrative predictors combining multiple information sources usually outperform sequence-based methods, these extra features are often unavailable, incomplete or difficult to obtain for most proteins thus limiting their scopes. Therefore, methods that accurately predict protein functions from sequences alone may be more general and applicable to most proteins that have not been extensively studied.

Since protein sequences can be regarded as a language in life, unsupervised pretraining with language models from natural language processing has recently been applied to protein sequence representation learning and has displayed promising results in downstream predictions including secondary structures, tertiary contacts, mutational effects, and protein binding sites [26-29]. Our previous work [29] has shown that sequence representations from pretrained language models can outperform manually-engineered evolutionary and structural features for binding site detection. Such results inspire us to develop a fast and accurate sequence-based protein function predictor that does not rely on any features constructed from protein domains, structures, biological networks or literature. Besides, network propagation approaches have been shown successful to predict protein functions in which existing knowledge is amplified by propagating an initial set of functional labels from experimentally characterized proteins through PPI networks [30]. S2F [22] further presents a label diffusion algorithm accounting for the overlapping communities of proteins with related functions. Therefore, it is promising to advance the performance of sequence-based function predictors by employing label diffusion over homology network built solely on sequence similarities.

In this study, we propose SPROF-GO, a Sequence-based alignment-free PROtein Function predictor, which leverages a pretrained protein language model to efficiently extract informative sequence embeddings and employs self-attention pooling to focus on important residues. Label diffusion algorithm is adopted to exploit the homology information and account for the overlapping communities of proteins with related functions. Besides, a hierarchical learning strategy is applied to produce consistent predictions and improve performance. SPROF-GO was shown to surpass state-of-the-art sequence-based and even network-based approaches by more than 14.5%, 27.3% and 10.1% in AUPR on the three sub-ontology test sets, respectively. Our method was further demonstrated to generalize well on non-homologous proteins and unseen species. Finally, visualization based on the attention mechanism indicated that SPROF-GO is able to capture sequence domains useful for function prediction. We suggest that our fast and accurate method could scale with the current fast-growing sequence databases, and provide useful information for biologists studying disease mechanism and chemists interested in targeted drug design.

## 2. Materials and methods

### 2.1 Datasets

We adopted the benchmark datasets proposed in [23], in which the training and test sets were collected following the standard protocol of CAFA. Specifically, the protein sequences were downloaded from UniProt [3], and the GO term annotations were extracted and combined from Swiss-Prot [31], GOA [32] and GO [6] in January 2020. Only experimental annotations with the following evidence codes were kept: IDA, IPI, EXP, IGI, IMP, IEP, IC or TA. The annotations were further up-propagated based on the “is-a” relation in the hierarchical structure of GO, and the root GO terms were omitted. Then, the training, validation and test sets were split according to the annotation time stamps. The training sets contain proteins annotated before January 2018, while the validation and test sets contain no-knowledge proteins annotated from January to December 2018 and from January 2019 to January 2020, respectively. In this study, we discarded sequences longer than 2,000 in the training sets and trimmed sequences to 5,000 in the validation and test sets due to the memory limit on GPU. Furthermore, to optimize the predicted accuracy, we only focused on the GO terms with enough training samples (≥ 50 sequences) in the training step, resulting in 790, 4766 and 667 classes for the MF, BP and CC sub-ontology. In the evaluation phase, we considered all terms to ensure fair comparison with other methods. **Table 1** shows the detailed statistics of the training, validation and test sets for the three domains of GO, as well as the HUMAN and MOUSE subsets used in our downstream analyses.

**Table 1.**
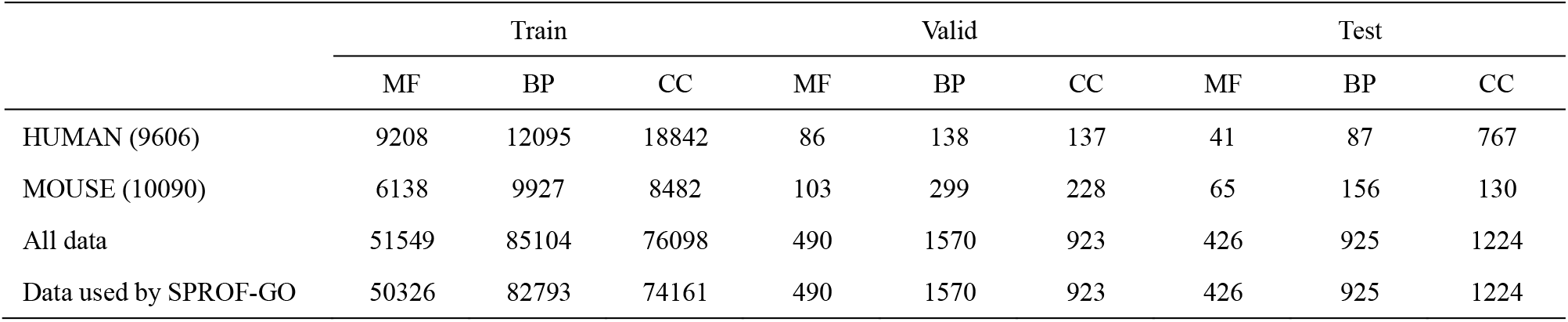
Statistics of the training, validation, and test sets used in this study for the three domains in GO

|  | Train |  |  | Valid |  |  | Test |  |  |
| --- | --- | --- | --- | --- | --- | --- | --- | --- | --- |
|  | MF | BP | CC | MF | BP | CC | MF | BP | CC |
| HUMAN (9606) | 9208 | 12095 | 18842 | 86 | 138 | 137 | 41 | 87 | 767 |
| MOUSE (10090) | 6138 | 9927 | 8482 | 103 | 299 | 228 | 65 | 156 | 130 |
| All data | 51549 | 85104 | 76098 | 490 | 1570 | 923 | 426 | 925 | 1224 |
| Data used by SPROF-GO | 50326 | 82793 | 74161 | 490 | 1570 | 923 | 426 | 925 | 1224 |

### 2.2 The architecture of SPROF-GO

The overall architecture of SPROF-GO is shown in **Figure 1**. First, the protein sequence is input to the pretrained protein language model to extract an initial sequence embedding matrix. Then, the embedding matrix is fed to two multilayer perceptrons (MLPs) parallelly to learn an attention vector and a more informative hidden embedding matrix. Finally, the hidden embeddings are weighted averaged among different sequence positions based on the attention scores to generate an embedding vector, which is input to the output MLP to predict the GO term probabilities. Additionally, a hierarchical learning strategy is applied to ensure the prediction to be consistent. This initial prediction is used during training to update the model parameters. In the test phase, the input sequence is further searched against the training set to build a sequence homology network. The initial prediction and the homology network are fed into the label diffusion algorithm, which outputs the final protein function prediction. Details of these modules are explained in the following sections.

**Figure 1.**
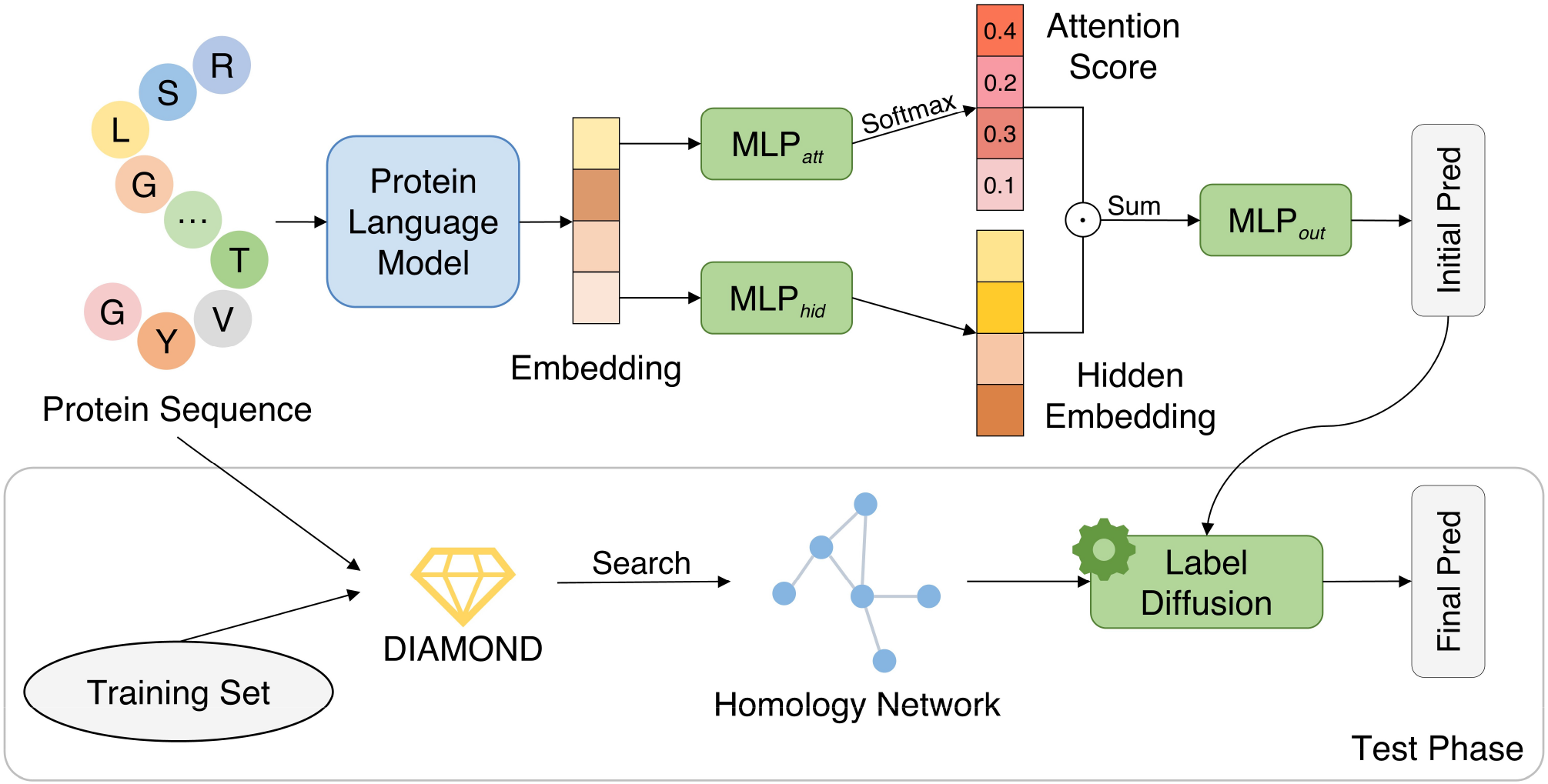
Overview of the SPROF-GO method. First, the protein sequence is input to the pretrained protein language model to extract the initial sequence embedding. Then, the embedding matrix is fed to two MLPs parallelly to learn an attention vector and a hidden embedding matrix. Finally, the hidden embeddings are weighted averaged among different sequence positions based on the attention scores, which is input to the output MLP to predict the GO term probabilities. This initial prediction is used during training to update the model parameters. In the test phase, the input sequence is further searched against the training set using DIAMOND to build a sequence homology network. The initial prediction and the homology network are fed into the label diffusion algorithm, which outputs the final protein function prediction.

### 2.2.1 Pretrained protein language model

SPROF-GO leverages the protein language model ProtT5-XL-U50 [27] (denoted as ProtTrans) for efficient feature extraction, thus bypassing the computationally intense sequence alignment to search for sequence domains or produce evolutionary profiles. ProtTrans is a transformer-based auto-encoder named T5 [33] pretrained in a self-supervised manner, essentially learning to predict masked amino acids. Concretely, the ProtTrans model contains 24 layers and 32 heads with 3B parameters, which was first trained on BFD [34] and then fine-tuned on UniRef50 [35]. The BERT’s denoising objective [36] was adopted to corrupt and reconstruct single tokens using a masking probability of 15% (details shown in **Supplementary Note 1**). We extracted the output from the last layer of the encoder part of ProtTrans as the initial sequence representation ***H***^(0)^ ∈ ℝ^n×1024^, with *n* denoting the sequence length and 1,024 being the feature dimension. Note that the inference cost of ProtTrans is really low, and the feature extraction process for our whole benchmark datasets (∼120,000 sequences, ∼500 amino acids on average) can be done in about 6 hours on an Nvidia GeForce RTX 3090 GPU. The feature values in the sequence representations from ProtTrans were further normalized to scores between 0 to 1 as follows:

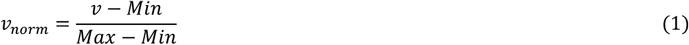

where *v* is the original feature value, and *Min* and *Max* are the smallest and biggest values of this feature type observed in the training set, respectively.

### Multilayer perceptron (MLP)

The sequence embedding matrix output from ProtTrans is fed to two MLPs parallelly to learn an attention vector and a hidden embedding matrix. MLP is a fully connected class of feedforward artificial neural network, which can be generally computed as follows:

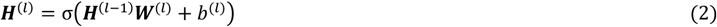

where 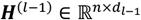 is the input of the *l*^th^ MLP layer; 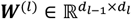 is the weight matrix; 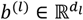 is the bias term; *σ* is the non-linear activation function; and 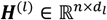 is the output of the l MLP layer. Between two layers of the MLP, we also add layer normalization [37] to stabilize the hidden state dynamics and dropout [38] to avoid overfitting.

### 2.2.3 Self-attention pooling

Many methods [16, 25] employ global mean pooling or max pooling to convert a residual-level embedding matrix into a protein-level embedding vector for subsequent function prediction, which might either dilute or lose the important features. Here we employ self-attention pooling to automatically focus on important residues as well as provide visualization and interpretability. We set the output dimension of the last layer in MLP_*att*_ to 1 and the activation function to softmax, so that the output of MLP_*att*_ is an attention vector ***A*** ∈ ℝ^n×1^. Let ***H***^(*L*)^ ∈ ℝ^*n*×*d*^ denote the hidden embedding output by MLP_*hid*_, then the self-attention pooling is calculated as follows:

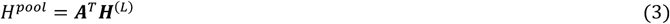

To jointly attend to information from different representation subspaces at different positions, multi-head attention is used in practice to produce *h* different attention vectors, perform the self-attention pooling in parallel and then concatenate them together. Finally, *H*^*pool*^ is input to MLP_*out*_ with a sigmoid function in the last layer to transform this embedding vector to a *K*-dimensional function prediction vector *H*^*out*^, where *K* is the number of GO terms that need to be predicted.

### 2.2.4 Hierarchical learning

Protein function prediction is a hierarchical multi-label classification problem, in which classes (go terms) are organized as a DAG, and every prediction must be consistent: the probability of a GO term must be equal to or greater than all of its child terms. Most methods (e.g., [22]) allow inconsistent predictions and require additional post-processing to ensure the consistency at inference time. Here we apply the hierarchical learning strategy proposed by [39] to produce consistent predictions and improve performance, which consists of two elements: (1) a max constraint module (MCM) built upon the network to guarantee consistent predictions inherently; (2) a loss function teaching the network when to exploit the predictions of the lower classes in the hierarchy for making predictions on the upper ones.

Specifically, let ***H*** be a *K* × *K* matrix obtained by stacking *K* rows of the prediction vector *H*^*out*^, and ***M*** be a *K* × *K* matrix such that ***M***_*ij*_ = 1 if the *j*^*th*^ GO term is a subclass of the *j*^*th*^ GO term, and ***M***_*ij*_ = 0 otherwise. Here, the subclasses of a target term include the child terms in the GO DAG and the target term itself, and only the “is-a” relation in GO is considered. Then, the prediction output by MCM is computed as:

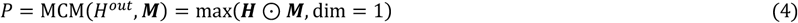

where ⊙ represents the element-wise product. In the validation and test phases, MCM sets the probability of a go term to the maximal probabilities of its subclasses, similar to the post-processing used by other methods. However, if the output of MCM is directly used for training with standard binary cross-entropy loss (BCELoss), the network may remain stuck in bad local optimal [39]. Thus, max constraint loss (MCLoss) is introduced to control when to exploit the predictive probabilities of the lower classes. Let *y* be the ground truth function annotation vector, 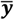 be a *K* x *K* matrix obtained by stacking *K* rows of *y*. Then the MCLoss is calculate as:

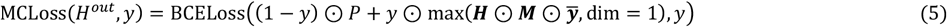

which means that the probability of a negative class should take the maximal probabilities of its subclasses, while the probability of a positive class should take the maximal probabilities of its positive subclasses.

### 2.2.5 Homology-based label diffusion

Proteins rarely perform their functions in isolation. Network propagation methods exploit the fact that groups of proteins connected in functional networks form communities that share similar functions [30]. However, when a protein has more than one function, it will belong to more than one functional group. Such proteins lying at the intersection of communities are generally more functionally similar compared to their neighbors, because they share more functional roles. Therefore, the propagation/diffusion of information between them should be higher. Here, we adopt the label diffusion algorithm proposed by [22] to explicitly model this overlapping community effect, in which we make three modifications: (1) we diffuse annotations over a homology network built solely on sequence similarities, instead of PPI networks in STRING; (2) we incorporate the ground truth annotations from the training set rather than only use test proteins for diffusion; (3) we employ DIAMOND [40] instead of BLAST [12] for similarity search and re-implement the algorithm with sparse matrix operation throughout, to accelerate the computation thus adapting to the large size of the training set.

Specifically, label diffusion is performed only in the test phase to further improve the initial function predictions. We use DIAMOND to search the whole test set against the training set to find training sequences similar to the test sequences, from which a homology network ***Q*** ∈ ℝ^*N*×*N*^ is built using the sequence identity for each pair of proteins (*N* is the number of hits in the training set plus the number of test proteins). Then, the weighted Jaccard similarity matrix is defined to measure how much a pair of proteins belong to the same community in network ***Q***:

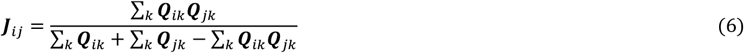

For a target go term *k*, we learn the *k*^*th*^ column of the final annotation matrix ***F*** (denoted as ***F***_*k*_) by minimizing the cost function *C*(***F***_*k*_):

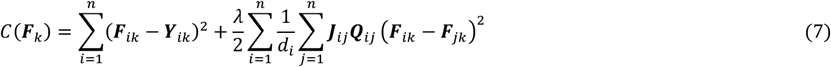

where the first term is to conserve the initial labels/predictions ***Y***_*ik*_, the second term accounts for the consistency of the labels/predictions of adjacent nodes in the network, ***J***_*ij*_***Q***_*ij*_ models the homology information and overlapping community effect, and *λ* is a regularization parameter. Note that 1⁄*d*_*i*_ is a normalization factor defined as:

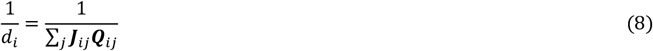

We define ***Q***\* and its Laplacian matrix ***L*** as follows:

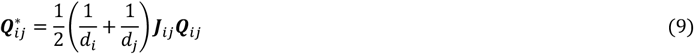

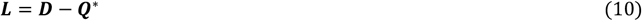

where ***D*** denotes the diagonal degree matrix of ***Q***\*. Then, the closed-form solution that minimizes *C*(***F***_*k*_) can be expressed as:

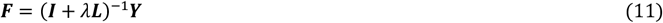

where ***I*** ∈ ℝ^*N*×*N*^ is an identity matrix, ***Y*** ∈ ℝ^*N*×*K*^is the concatenation of the labels of the training set and the initial predictions of the test set, and ***F*** ∈ ℝ^*N*x*K*^ is the updated annotations for the training samples and the test proteins, from which we retrieve our final predictions for the test sequences. Moreover, as proven in [22], since our initial predictions are consistent with the GO structure, our final predictions output by label diffusion will also be consistent.

### 2.3 Implementation and evaluation

We trained our models to predict GO terms separately for MF, BP and CC ontology. For each training set of the sub-ontology, we trained five models using five different random seeds, and their average performance on the validation set was used to choose the best feature combination and optimize all hyperparameters through grid search (**Supplementary Table S1**). In the test phase, all five trained models were used to make predictions, which were then averaged as the assembled prediction of SPROF-GO. Specifically, we employed a two-layer fully connected architecture for the three MLPs in SPROF-GO with the following set of hyperparameters: hidden units of 256, attention heads (*h*) of 8, dropout rate of 0.1, and batch size of 20. The label diffusion regularization parameter *λ* was simply set to 1. We utilized the Adam optimizer [41] with *β*_*1*_ = 0.9, *β*_*2*_ =0.99, weight decay of 10^−5^ and learning rate of 2 × 10^−4^ for model optimization. We implemented the proposed method with Pytorch 1.13.0 [42]. The training process for one model lasted at most 30 epochs and we performed early stopping with patience of 4 epochs based on the validation performance, which took ∼40 minutes for MF and CC ontology, and ∼2 hours for BP ontology on an Nvidia GeForce RTX 3090 GPU. During the test phase, it took ∼2 minutes to make predictions for all proteins in the three sub-ontology test sets (∼2,500 sequences).

Similar to the previous studies [14, 23], we used F_max_ and AUPR (area under the precision-recall curve) to evaluate the predictive performance, whose detailed definitions are given in **Supplementary Note 2**. F_max_ is the maximum protein-centric F-measure computed over all prediction thresholds, which is a major evaluation metrics in CAFA. AUPR is also a suitable measure for highly unbalanced dataset since it emphasizes more on the minority class [43, 44].

## 3. Results

The overview of SPROF-GO is shown in **Figure 1**. For a given protein sequence, the pretrained language model is employed to extract a sequence embedding matrix, which is fed to the self-attention pooling module to generate a protein-level representation for the final output layer. Besides, hierarchical learning is applied to ensure consistent prediction. In the test phase, the prediction is further advanced by exploiting the homology information through label diffusion. The Results section is organized as follows. First, we demonstrated the superiority of the feature from pretrained language model over other widely used features. Second, we conducted ablation study on several techniques used in SPROF-GO. Third, we compared SPROF-GO with other state-of-the-art methods. Fourth, we evaluated SPROF-GO on non-homologous proteins and unseen species to verify its robustness. Lastly, we interpreted the decision mechanism of SPROF-GO by visualizing the attention scores.

### 3.1 Feature from pretrained language model is informative for protein function prediction

We evaluated SPROF-GO on the test sets of the three domains in GO (described in **Table 1**) by F_max_ and AUPR. As shown in **Table 2**, SPROF-GO achieved F_max_ of 0.647, 0.335, and 0.725, as well as AUPR of 0.622, 0.247, and 0.765 on the MF, BP, and CC test sets, respectively. To demonstrate the effectiveness of the language model (ProtTrans) representation employed by SPROF-GO, we conducted feature ablation experiments to compare ProtTrans with other popular features in this field. When adopting the one-hot encoding of amino acid types, the model showed poor performance with AUPR of 0.475, 0.187, and 0.705 on the three sub-ontology test sets, and the performance would further degrade largely when removing the label diffusion module (AUPR of 0.177, 0.130, and 0.582). This suggested that the primary sequences alone are insufficient to characterize protein functions, while sequence homology information is still a valuable source for function inference. We also investigated the widely used [11, 19, 23, 25] InterPro feature generated by InterProScan [13] through sequence alignment, which is a binary protein-level vector indicating the existences of protein domains and families. As shown in **Table 2**, the model using InterPro obtained AUPR of 0.594, 0.203, and 0.730, surpassing the one using one-hot encoding, which is reasonable since domains often form functional units, such as the calcium-binding EF hand domain of calmodulin. However, the sequence feature by ProtTrans outperformed one-hot, InterPro or the combination of these two features by large margins. Note that the generation of the ProtTrans feature is also much more efficient than that of InterPro since no database searches are needed. Moreover, further integrating one-hot and InterPro features to ProtTrans was redundant and could not attain any further improvements, suggesting that the ProtTrans language model may have potentially captured the protein sequence, domain and family knowledge informative for function prediction.

**Table 2.** The predictive performance on the test sets of the three domains in GO using different features.

| Feature | F<sub>max</sub> |  |  | AUPR |  |  |
| --- | --- | --- | --- | --- | --- | --- |
|  | MF | BP | CC | MF | BP | CC |
| One-hot | 0.555 | 0.291 | 0.680 | 0.475 | 0.187 | 0.705 |
| InterPro | 0.631 | 0.300 | 0.687 | 0.594 | 0.203 | 0.730 |
| One-hot + InterPro | 0.634 | 0.299 | 0.689 | 0.589 | 0.203 | 0.732 |
| ProtTrans (SPROF-GO) | **0.647** | **0.335** | **0.725** | **0.622** | **0.247** | **0.765** |
| ProtTrans + one-hot + InterPro | 0.645 | 0.329 | 0.722 | 0.620 | 0.239 | 0.750 |

*Note:* Bold fonts indicate the best results.

### 3.2 Model ablation study

To investigate the impacts of the self-attention pooling, hierarchical learning, homology-based label diffusion, and model assembly techniques applied by SPROF-GO, we removed one of the four components at a time and then re-trained the model using the same sequence feature. As shown in **Table 3**, the removal of the assembly strategy caused the largest average AUPR drop (0.017) on the three test sets, while the removals of the attention pooling (using mean pooling instead), hierarchical learning (using post-processing instead), and label diffusion caused average AUPR drops of 0.009, 0.008, and 0.010, respectively. Note that since SPROF-GO is supported by several techniques, removal of a single component seemed to have minor influence on the overall performance. Moreover, some components may have significant benefits on one ontology, but have little impacts on the others. For example, the removal of label diffusion caused the largest AUPR drop of 0.022 on the MF test set, while it had no impact on the AUPR of the CC test set. The removal of attention or assembly caused the largest AUPR drops of 0.012 on the BP test set, and the removal of assembly caused the largest AUPR drop of 0.022 on the CC test set. Here, we also report the performance of a baseline method that doesn’t use any of the above-mentioned techniques (SPROF-GO_base_). SPROF-GO outperformed this baseline significantly with improvements of 0.064, 0.028, and 0.033 on F_max_, and 0.059, 0.038, and 0.032 on AUPR in the three test sets, further indicating the advantages of the four modules in SPROF-GO.

**Table 3.** Ablation study on different techniques used by SPROF-GO on the test sets of the three domains in GO.

| Method | F<sub>max</sub> |  |  | AUPR |  |  |
| --- | --- | --- | --- | --- | --- | --- |
|  | MF | BP | CC | MF | BP | CC |
| SPROF-GO<sub>base</sub> | 0.583 | 0.307 | 0.692 | 0.563 | 0.209 | 0.733 |
| SPROF-GO w/o attention | 0.628 | 0.328 | 0.721 | 0.613 | 0.235 | 0.760 |
| SPROF-GO w/o hierarchical learning | 0.638 | 0.333 | 0.723 | 0.612 | 0.238 | 0.761 |
| SPROF-GO w/o label diffusion | 0.633 | 0.328 | 0.721 | 0.600 | 0.240 | **0.765** |
| SPROF-GO w/o assembly | 0.646 | 0.327 | 0.715 | 0.605 | 0.235 | 0.743 |
| SPROF-GO | **0.647** | **0.335** | **0.725** | **0.622** | **0.247** | **0.765** |

*Note:* SPROF-GO<sub>base</sub> denotes the baseline method that doesn't use any of the above-mentioned techniques. Bold fonts indicate the best results.

### 3.3 Comparison with state-of-the-art methods

We compared SPROF-GO with four sequence-based (BLAST-KNN, LR-InterPro, DeepGOCNN, and DeepGOPlus) and two network-based (Net-KNN and DeepGraphGO) predictors on the test sets of the three domains in GO. The baseline method (SPROF-GO_base_) that utilizes ProtTrans and MLP with mean pooling is also considered here. The implementation details of these competing methods are introduced in **Supplementary Note 3**. As reported in **Table 4**, GO terms in the BP ontology seem harder to predict for all methods, which may be due to the large number of terms and the deep and complex structure of the BP DAG. Howsoever, SPROF-GO outperformed all other sequence-based and even network-based methods on F_max_ and AUPR in all three domains. Undoubtedly, SPROF-GO substantially surpassed the sequence-based method DeepGOPlus by 56.3%, 128.7% and 28.6% on AUPR in the three test sets, respectively. This indicated that representing protein sequence simply by one-hot encoding followed by CNN, and mining homology information simply using k-nearest neighbors are not enough to capture the most helpful information for function prediction. Interestingly, though our method is a sequence-based predictor, it outperformed the state-of-the-art network-based method DeepGraphGO by 3.9%, 2.4% and 4.8% on F_max_, and 14.5%, 27.3% and 10.1% on AUPR. This is expected because the sequence representation from the pretrained language model used by SPROF-GO is more informative and powerful than the handcrafted domain and family features employed by DeepGraphGO (shown in **section 3.1**). Another reason may be that the network information also brought noises since the protein-protein associations in STRING are not always from experiments. In addition, the label diffusion in SPROF-GO could further boost the predictive quality by exploiting the homology information and overlapping community effect. On the other hand, our method is also computationally efficient. Empirically, it takes ∼7 minutes to extract features and make predictions on the three ontologies for 1,000 proteins with 500 residues on average using SPROF-GO on an Nvidia GeForce RTX 3090 GPU. However, DeepGraphGO can only predict less than 5 sequences in the same time since the generation of the InterPro feature requires expensive database searches.

**Table 4.** Performance comparison of SPROF-GO with state-of-the-art methods on the test sets of the three domains in GO.

| Method | F<sub>max</sub> |  |  | AUPR |  |  |
| --- | --- | --- | --- | --- | --- | --- |
|  | MF | BP | CC | MF | BP | CC |
| BLAST-KNN | 0.590 | 0.274 | 0.650 | 0.455 | 0.113 | 0.570 |
| LR-InterPro | 0.617 | 0.278 | 0.661 | 0.530 | 0.144 | 0.672 |
| Net-KNN | 0.426 | 0.305 | 0.667 | 0.276 | 0.157 | 0.641 |
| DeepGOCNN | 0.434 | 0.248 | 0.632 | 0.306 | 0.101 | 0.573 |
| DeepGOPlus | 0.593 | 0.290 | 0.672 | 0.398 | 0.108 | 0.595 |
| DeepGraphGO | <u>0.623</u> | <u>0.327</u> | <u>0.692</u> | 0.543 | 0.194 | 0.695 |
| SPROF-GO<sub>base</sub> | 0.583 | 0.307 | <u>0.692</u> | <u>0.563</u> | <u>0.209</u> | <u>0.733</u> |
| SPROF-GO | **0.647** | **0.335** | **0.725** | **0.622** | **0.247** | **0.765** |

*Note:* SPROF-GO<sub>base</sub> denotes the baseline method that employs ProtTrans and MLP with mean pooling. Bold and underlined fonts indicate the best and second-best results, respectively.

### 3.4 Generalization on non-homologous proteins and unseen species

To examine the generalization ability of our method for non-homologous proteins, we compared SPROF-GO with other competing methods on *difficult* proteins within the test sets, which are defined by CAFA2 [8] as the test proteins with sequence identity <60% to the training set. The numbers of *difficult* proteins in the MF, BP, and CC test sets are 303, 649, and 437, respectively. As shown in **Table 5**, almost all methods showed performance drops in different degrees compared to the results on the original test sets (**Table 4**). However, SPROF-GO still outperformed all other methods on F_max_ and AUPR in all three domains. Specifically, SPROF-GO maintained similar performance in the MF/BP ontology, with AUPR of 0.622/0.247 on the full test set and 0.617/0.256 on the subset of *difficult* proteins. By comparison, the AUPR of DeepGraphGO in MF and BP decreased from 0.543 to 0.508 and 0.194 to 0.184, respectively. As for the CC ontology, AUPR decreased by 12.7% for DeepGraphGO but only 7.5% for SPROF-GO on *difficult* proteins. Interestingly, we found that the label diffusion module could still bring improvements in this scenario by mining annotations from dissimilar sequences, and its removal caused AUPR drops of 0.020, 0.011 and 0.003 on the MF, BP, and CC test sets. These results suggested that our method can generalize well on non-homologous proteins, rather than just remember the functions of similar sequences in the training set, thus making it a robust and reliable method for function prediction of novel sequences.

**Table 5.** Performance comparison of SPROF-GO with state-of-the-art methods on *difficult* proteins within the test sets of the three domains in GO.

| Method | $F_{\max}$ | | | AUPR | | |
| --- | --- | --- | --- | --- | --- | --- |
|  | MF | BP | CC | MF | BP | CC |
| BLAST-KNN | 0.534 | 0.274 | 0.521 | 0.377 | 0.114 | 0.354 |
| LR-InterPro | 0.589 | 0.275 | 0.613 | 0.493 | 0.148 | 0.591 |
| Net-KNN | 0.404 | 0.292 | 0.595 | 0.230 | 0.142 | 0.560 |
| DeepGOCNN | 0.406 | 0.243 | 0.578 | 0.246 | 0.091 | 0.478 |
| DeepGOPlus | 0.564 | 0.292 | 0.602 | 0.326 | 0.108 | 0.454 |
| DeepGraphGO | <u>0.598</u> | <u>0.322</u> | <u>0.625</u> | <u>0.508</u> | <u>0.184</u> | <u>0.607</u> |
| SPROF-GO | **0.630** | **0.339** | **0.682** | **0.617** | **0.256** | **0.708** |

Note: Bold and underlined fonts indicate the best and second-best results, respectively.

We also investigated the performance of SPROF-GO and other methods over proteins of HUMAN and MOUSE within the test sets (details shown in **Table 1**), and SPROF-GO again outperformed all competing methods in all twelve settings except one (**Supplementary Table S2**). More importantly, to explore the generalization ability of our method for unseen species, we further evaluated SPROF-GO and the second-best method DeepGraphGO on the HUMAN and MOUSE proteins when trained with proteins of all species except the target species (denoted by w/o species). As shown in **Figure 2**, the AUPR of both methods decreased in most cases when excluding the training data of the target species. However, SPROF-GO_w/o species_ still surpassed DeepGraphGO_w/o species_ in all three domains for the HUMAN and MOUSE proteins. Moreover, even without training data from the target species, SPROF-GO_w/o species_ outperformed DeepGraphGO using the whole training set in all six settings except one. For example, SPROF-GO_w/o HUMAN_ achieved AUPR of 0.735 for CC ontology on the HUMAN proteins, exceeding the one by DeepGraphGO using the full training set (0.642). Detailed evaluation results are shown in **Supplementary Table S3** and **S4**, where the performance using only the target species for training is also included. These results suggested that our method is robust and can also generalize on proteins of unseen species in the training set, thus having the potential to predict functions for newly sequenced organisms.

**Figure 2.**
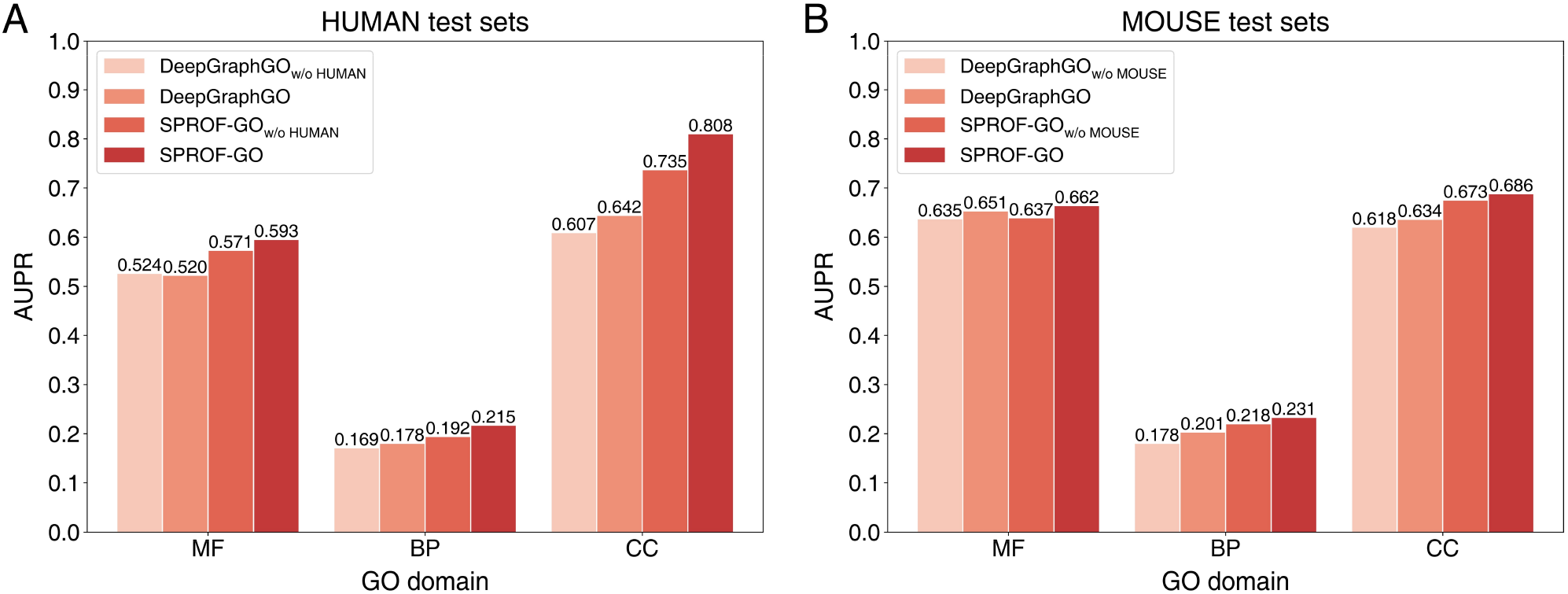
Performance comparison of SPROF-GO and DeepGraphGO on the HUMAN (**A**) and MOUSE (**B**) proteins within the test sets when trained with proteins of all species or all species except the target species (denoted by w/o HUMAN and w/o MOUSE).

### 3.5 Model interpretation by attention visualization

What did SPROF-GO learn? Did the network reason solely by comparing the test proteins with the training samples or did it learn the underlying chemical principles of protein functioning? To better illustrate the decision mechanism of SPROF-GO, we selected two examples (UniProt ID: Q6DR03 and Q9LIB5) in the MF ontology test set and extracted their residue-level attention scores from the self-attention pooling module in SPROF-GO. We averaged the scores from different attention heads and different assembled models as the final attention scores. Besides, we applied InterProScan to search for functional domains in these two sequences. As shown in **Figure 3**, Q6DR03 contains a DHHC domain of palmitoyltransferases [45] in sequence positions of 140 to 277 (in blue), where the attention scores (in red) are also higher. This protein was annotated hierarchically with five GO terms down to “S-acyltransferase activity”, in which SPROF-GO correctly predicted four terms but missed one specific term, leading to the F-measure of 0.889. Another case Q9LIB5 contains a GATA-type zinc finger domain [46] in sequence positions of 38 to 93, where the attention scores by SPROF-GO are also higher. The presence of the zinc finger domain associates this protein with several functions involving DNA-binding, in which SPROF-GO correctly predicted all seven terms, leading to an F-measure of 1.000. The attention visualization for these cases suggested that our method could successfully identify and pay more attention to the functional domains in sequences, and thus correctly predicted the correlated functions of the proteins.

**Figure 3.**
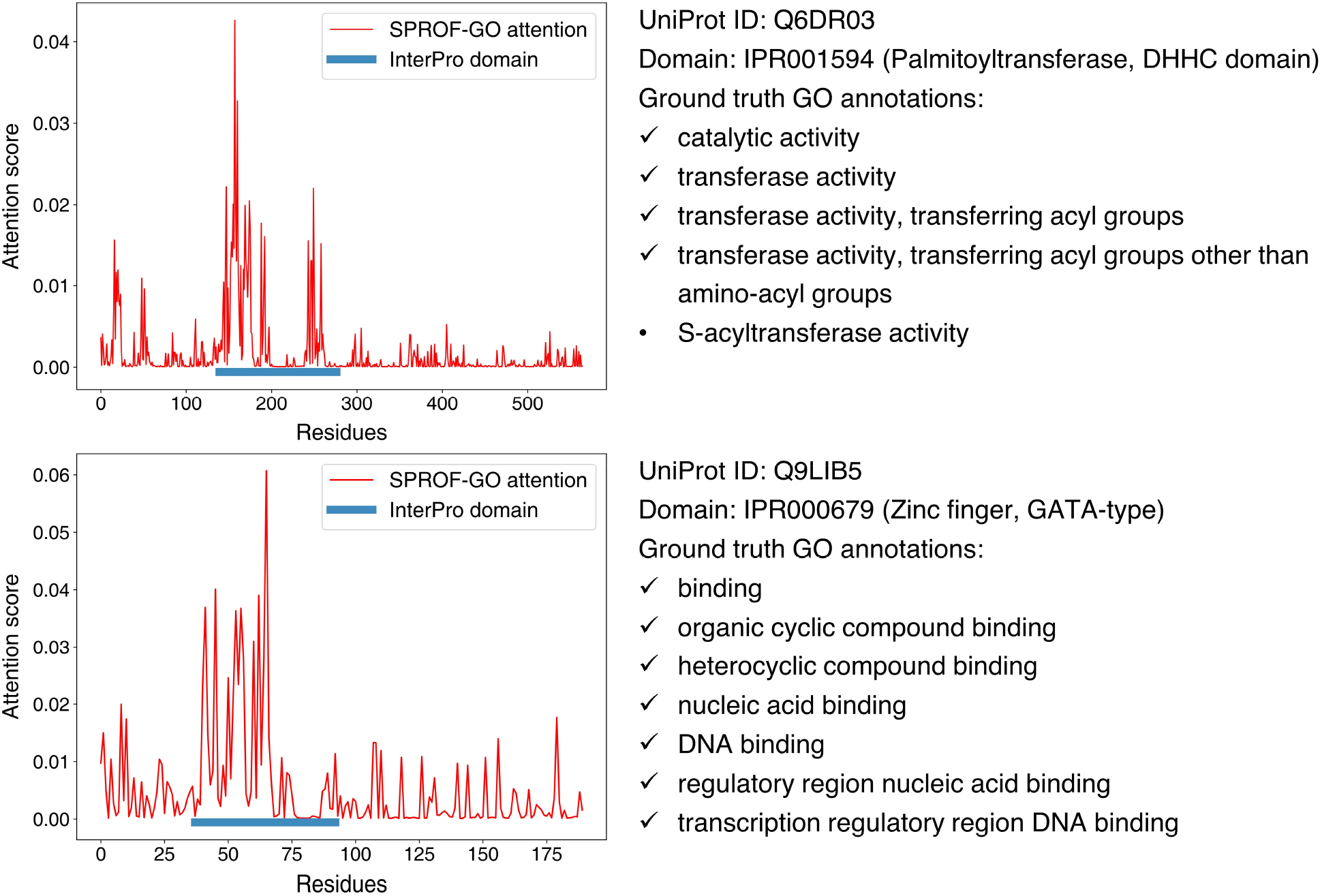
Visualization of two examples (Q6DR03 and Q9LIB5) in the MF ontology test set. The left panels show the attention scores by SPROF-GO at different sequence positions (in red), as well as the locations of the domains found by InterProScan (in blue). The right panels show the information of the test proteins, including UniProt ID, InterPro domain ID, domain name, and ground truth function annotation. The GO terms correctly identified by SPROF-GO are marked with ticks, and the root GO term (GO:0003674 molecular function) is omitted.

## 4. Discussion

Protein function prediction benefits disease mechanism elucidation and drug target discovery. Existing sequence-based methods mostly suffer from low predictive accuracy or high computational cost. Although many methods have integrated protein structures, biological networks or literature information to improve performance, these extra features are often unavailable. Here, we propose a Sequence-based PROtein Function predictor SPROF-GO, which has the following five notable features: (1) SPROF-GO leverages a pretrained language model to efficiently extract informative sequence embeddings, thus bypassing expensive database searches; (2) The self-attention pooling is employed to capture sequence domains useful for function prediction and provide interpretability; (3) SPROF-GO applies a hierarchical learning strategy to produce consistent predictions and improve performance; (4) The label diffusion algorithm is adopted to exploit the homology information and overlapping community effect; (5) SPROF-GO is accurate and robust, with better performance than state-of-the-art sequence-based and even network-based approaches, and great generalization ability on non-homologous proteins and unseen species.

However, there is still room for further improvements on SPROF-GO. First, applying GNN [47, 48] on predicted protein structures from sequences by AlphaFold2 [49] or ESMFold [50] may yield better performance. Second, contrastive learning [51] could be applied on the PPI networks only in the training phase to maximize the function similarities between network neighbors, thus reflecting the guilt-by-association principle. Third, knowledge graph techniques [52] could also be explored in this problem to incorporate drug and disease information. Fourth, SPROF-GO currently only considers around 6,200 GO terms that have ≥ 50 training samples, since we found that predicting all terms would lead to performance degrade. How to effectively learn the terms with scarce training data remains a challenging and interesting task to solve in our future. We suggest that our fast and accurate method could scale with the current fast-growing sequence databases, and provide useful information for biologists studying disease mechanism and chemists interested in targeted drug design.

## Supplementary Information

### Supplementary Note 1

SPROF-GO leverages the protein language model ProtT5-XL-U50 [1] (denoted as ProtTrans) for feature extraction, which is a transformer-based auto-encoder named T5 [2] pretrained in a self-supervised manner. Concretely, the ProtTrans model was first trained on BFD (Big Fantastic Database) [3] for 1.2M steps with 1024 Google TPUs and then fine-tuned on UniRef50 [4] for 991k steps with 256 TPUs. BFD contains 2,122 million proteins with 393 billion amino acids, while UniRef50 contains 45 million proteins with 14 billion amino acids. Non-generic or unresolved amino acids ([BOUZ]) were all mapped to X. Contrary to the original T5 model which masks spans of multiple tokens, ProtTrans adopts the BERT’s denoising objective [5] to corrupt and reconstruct single tokens using a masking probability of 15%. As for the network structure, ProtTrans uses the original transformer architecture proposed for sequence translation, which consists of an encoder that projects a source language to an embedding space and a decoder that generates a translation to a target language based on the encoder’s embedding. Specifically, the model contains 24 layers and 32 heads with hidden dimension of 1024. The model contains 3B parameters and was trained using 8-way model parallelism with batch size of 4 for BFD and 8 for UniRef50, respectively. The AdaFactor optimizer with weight decay of 0 and learning rate of 0.01 was utilized for model optimization. The dropout rate was set to 0.1 to avoid overfitting. The pretrained ProtTrans model is now available through https://github.com/agemagician/ProtTrans.

### Supplementary Note 2

We evaluated the predictive performance for the three domains in GO (MF, BP, and CC) independently using F_max_ (maximum protein-centric F-measure) and AUPR (area under the precision-recall curve). F_max_ is a major evaluation metrics in CAFA [6], which is computed as follows:

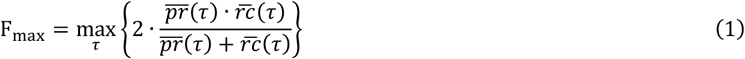

where 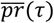 and 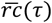 denote the average precision and recall at threshold *τ* respectively, which are defined as follows:

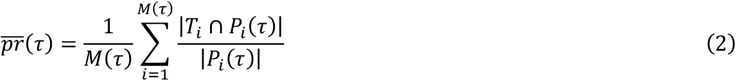

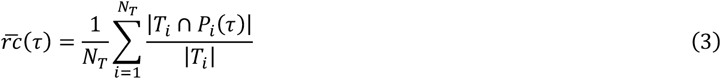

where *M*(*τ*) denotes the number of proteins predicted with at least one GO term at threshold *τ*; *N*_*T*_ denotes the total number of proteins; *T*_*i*_ denotes the true annotation set for protein *i*; *P*_*i*_(*τ*) denotes the predicted annotation set for protein *i* at threshold *τ*; and |·| is the set cardinality operation. We used 101 thresholds from 0 to 1 with an interval of 0.01 to compute the above measurement. In addition, we also reported AUPR and used it for feature and hyperparameter selections as it is sensitive and informative and it emphasizes more on the minority class in unbalanced classification task [7, 8].

### Supplementary Note 3

We compared SPROF-GO with four sequence-based (BLAST-KNN, LR-InterPro, DeepGOCNN, and DeepGOPlus [9]) and two network-based (Net-KNN and DeepGraphGO [10]) predictors on the test sets of the three domains in GO. The baseline method (SPROF-GO_base_) that utilizes ProtTrans and employs MLP with mean pooling is also considered here. The evaluation results of these methods on the test sets are directly obtained from [10] (except for SPROF-GO_base_). Details of these competing methods are introduced below.

### BLAST-KNN

The idea of BLAST-KNN is that similar proteins may have similar functions. For a test protein *p*_*i*_, BLAST [11] is run with an e-value cut-off of 0.001 to search *p*_*i*_ against all proteins in the training set to obtain its homologous protein set *H*_*i*_. Then the probability score between *p*_*i*_ and a GO term *G*_*j*_ is computed as follows:

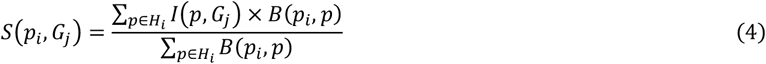

where *B*(*p*_*i*_, *p*) is the bit-score between *p*_*i*_ and *p*, and *I*(*p, G*_*j*_) is a binary indicator which equals to 1 if protein *p* has the function *G*_*j*_, or equals to 0 otherwise.

### LR-InterPro

The InterPro database [12] combines 14 different protein domain/family databases, including Pfam [13], CATH-Gene3D [14], CCD [15] and so on, covering a large number of domains, families, and motifs in sequences. The InterPro feature is generated by InterProScan [16], which is a binary protein-level feature vector indicating the existences of protein domains and families. This vector is used as an instance to train a logistic regression (LR) classifier for each GO term.

### DeepGOCNN and DeepGOPlus

DeepGOPlus is a sequence-based method consisting of two components, a deep convolutional neural network (CNN) named DeepGOCNN and a k-nearest neighbor method called DiamondScore. The DeepGOCNN module scans the sequence for motifs which are predictive for protein functions, and the DiamondScore exploits the homology information as in BLAST-KNN, except that the protein similarity is computed by DIAMOND [17].

### Net-KNN

The predictive score by Net-KNN is similar to that by BLAST-KNN, except that the similarities between proteins are now replaced by the weights in the protein-protein interaction (PPI) network from STRING [18]:

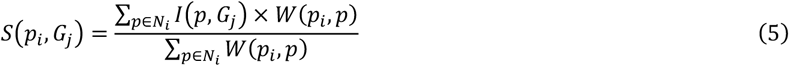

where *N*_*i*_ is the neighbor nodes (proteins) of *p*_*i*_ in the PPI network, and *W*(*p*_*i*_, *p*) is the weight of the edge between *p*_*i*_ and *p*. If the test protein *p*_*i*_ is not in the network, the prediction score of the most similar protein to *p*_*i*_ in the network is used as the prediction for *p*_*i*_.

### DeepGraphGO

DeepGraphGO is a network-based method which makes the most of both protein sequence domain and high-order protein network information via multispecies GNN strategy. DeepGraphGO has two inputs: (i) a graph (PPI network) with *N* nodes (proteins), which is formatted as a weighted adjacency matrix *A* ∈ ℝ^*N*×*N*^ (edge weights range between 0 and 1). (ii) *N* binary node feature vectors (InterPro). The prediction procedure has three steps: (i) Input MLP layer: the binary feature vector of each protein is transformed into a dense hidden vector (initial representation). (ii) Graph convolutional network (GCN): it captures the high-order information through graph edges to update the representation of each node (protein) using those of neighboring nodes. (iii) Output MLP layer: it predicts scores of GO terms for each protein.

### SPROF-GO_base_

SPROF-GO_base_ is a baseline method that utilizes the same ProtTrans feature as SPROF-GO and employs a 2-layer MLP with mean pooling. We post-process the raw outputs from SPROF-GO_base_ to ensure consistent predictions. Neither label diffusion nor model assembly is applied.

**Table S1.** The performance of SPROF-GO on the validation and test sets using different hyperparameters.

| Hyperparameter | Value | Validation<br>AUPR* | F<sub>max</sub> |  |  | AUPR |  |  |
| --- | --- | --- | --- | --- | --- | --- | --- | --- |
|  |  |  | MF | BP | CC | MF | BP | CC |
| Layers | 1 | 0.489 | 0.633 | 0.329 | 0.724 | 0.611 | 0.244 | 0.761 |
|  | **2** | **0.499** | 0.647 | 0.335 | 0.725 | 0.622 | 0.247 | 0.765 |
|  | 3 | 0.496 | 0.642 | 0.335 | 0.725 | 0.618 | 0.243 | 0.766 |
| Hidden units | 128 | 0.498 | 0.637 | 0.340 | 0.723 | 0.613 | 0.244 | 0.763 |
|  | **256** | **0.499** | 0.647 | 0.335 | 0.725 | 0.622 | 0.247 | 0.765 |
|  | 512 | 0.494 | 0.631 | 0.335 | 0.727 | 0.618 | 0.250 | 0.759 |
| Attention heads | 4 | 0.496 | 0.635 | 0.333 | 0.726 | 0.616 | 0.242 | 0.771 |
|  | **8** | **0.499** | 0.647 | 0.335 | 0.725 | 0.622 | 0.247 | 0.765 |
|  | 16 | 0.496 | 0.642 | 0.338 | 0.724 | 0.616 | 0.251 | 0.762 |
| Dropout | **0.1** | **0.499** | 0.647 | 0.335 | 0.725 | 0.622 | 0.247 | 0.765 |
|  | 0.2 | 0.495 | 0.650 | 0.332 | 0.727 | 0.616 | 0.244 | 0.766 |
|  | 0.3 | 0.498 | 0.644 | 0.331 | 0.725 | 0.613 | 0.242 | 0.763 |
| Learning rate | 1e-4 | 0.493 | 0.641 | 0.335 | 0.720 | 0.616 | 0.244 | 0.760 |
|  | **2e-4** | **0.499** | 0.647 | 0.335 | 0.725 | 0.622 | 0.247 | 0.765 |
|  | 5e-4 | 0.492 | 0.645 | 0.336 | 0.724 | 0.623 | 0.241 | 0.761 |

*Note:* For each row, the hyperparameters that are not shown are the same as the final SPROF-GO model. Validation AUPR\* denotes the average AUPR of the three domains of GO on five runs using different random seeds. Bold fonts indicate the best hyperparameters and the corresponding validation AUPR.

**Table S2.** Performance comparison of SPROF-GO with state-of-the-art methods on HUMAN and MOUSE proteins within the test sets of the three domains in GO.

| Method | F<sub>max</sub> |  |  | AUPR |  |  |
| --- | --- | --- | --- | --- | --- | --- |
|  | MF | BP | CC | MF | BP | CC |
| HUMAN (9606) |  |  |  |  |  |  |
| BLAST-KNN | 0.471 | 0.241 | 0.555 | 0.296 | 0.074 | 0.384 |
| LR-InterPro | 0.593 | 0.282 | 0.650 | 0.496 | 0.138 | 0.603 |
| Net-KNN | 0.485 | 0.261 | 0.615 | 0.358 | 0.143 | 0.620 |
| DeepGOCNN | 0.468 | 0.263 | 0.594 | 0.327 | 0.114 | 0.552 |
| DeepGOPlus | 0.501 | 0.277 | 0.625 | 0.246 | 0.088 | 0.479 |
| DeepGraphGO | <u>0.633</u> | **0.320** | <u>0.655</u> | <u>0.520</u> | <u>0.178</u> | <u>0.642</u> |
| SPROF-GO | **0.647** | <u>0.311</u> | **0.750** | **0.593** | **0.215** | **0.808** |
| MOUSE (10090) |  |  |  |  |  |  |
| BLAST-KNN | <u>0.681</u> | 0.289 | 0.593 | 0.593 | 0.105 | 0.441 |
| LR-InterPro | 0.628 | 0.312 | 0.592 | 0.625 | 0.175 | 0.569 |
| Net-KNN | 0.420 | 0.302 | 0.588 | 0.319 | 0.167 | 0.569 |
| DeepGOCNN | 0.475 | 0.258 | 0.574 | 0.405 | 0.129 | 0.495 |
| DeepGOPlus | 0.634 | 0.306 | 0.598 | 0.550 | 0.132 | 0.488 |
| DeepGraphGO | 0.650 | <u>0.329</u> | <u>0.638</u> | <u>0.651</u> | <u>0.201</u> | <u>0.634</u> |
| SPROF-GO | **0.686** | **0.332** | **0.676** | **0.662** | **0.231** | **0.686** |

*Note:* Bold and underlined fonts indicate the best and second-best results, respectively.

**Table S3.**
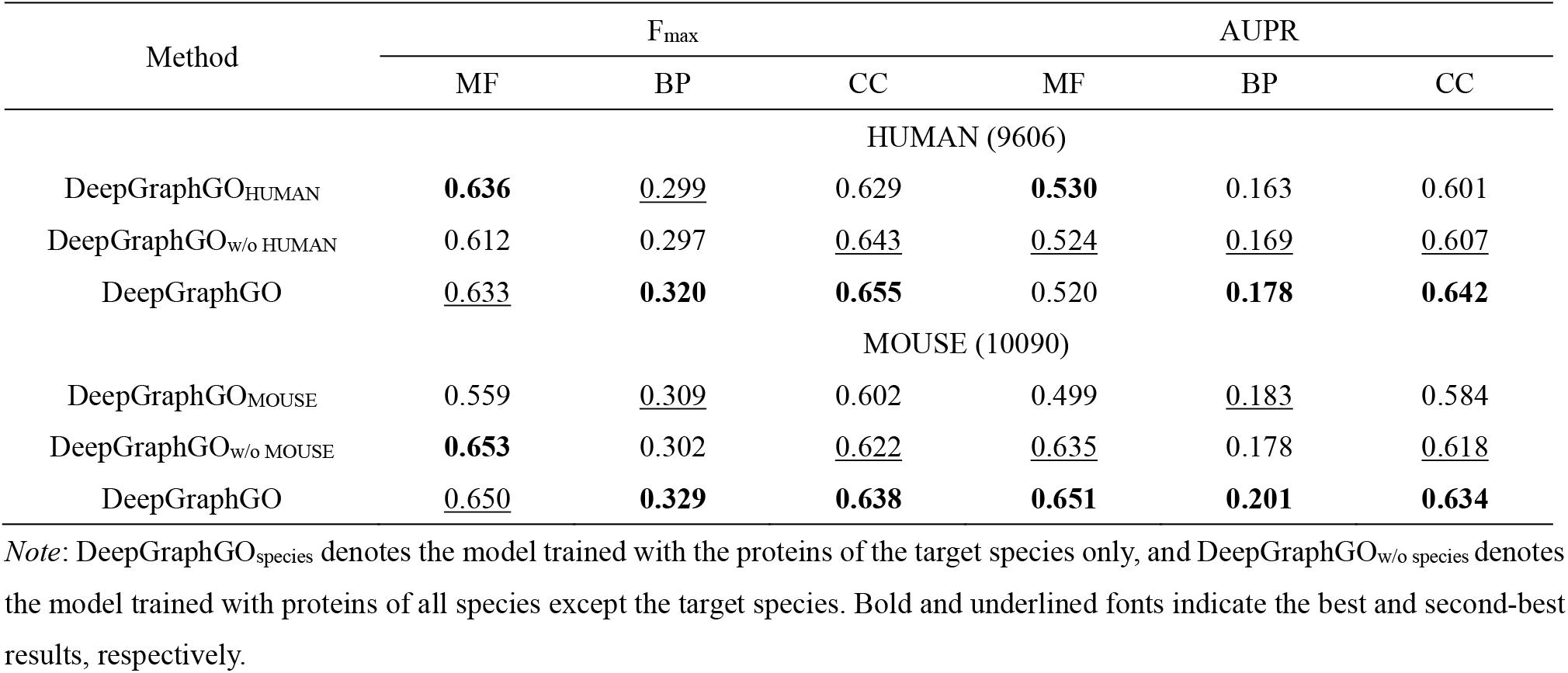
Performance comparison of DeepGraphGO and the two variants on the HUMAN and MOUSE proteins within the test sets of the three domains in GO.

**Table S4.** Performance comparison of SPROF-GO and the two variants on the HUMAN and MOUSE proteins within the test sets of the three domains in GO.

| Method | F<sub>max</sub> |  |  | AUPR |  |  |
| --- | --- | --- | --- | --- | --- | --- |
|  | MF | BP | CC | MF | BP | CC |
| HUMAN (9606) |  |  |  |  |  |  |
| SPROF-GO<sub>HUMAN</sub> | <u>0.634</u> | **0.325** | **0.765** | <u>0.572</u> | **0.219** | <u>0.790</u> |
| SPROF-GO<sub>w/o HUMAN</sub> | 0.628 | 0.289 | 0.679 | 0.571 | 0.192 | 0.735 |
| SPROF-GO | **0.647** | <u>0.311</u> | <u>0.750</u> | **0.593** | <u>0.215</u> | **0.808** |
| MOUSE (10090) |  |  |  |  |  |  |
| SPROF-GO<sub>MOUSE</sub> | **0.703** | 0.321 | <u>0.673</u> | <u>0.647</u> | <u>0.223</u> | 0.650 |
| SPROF-GO<sub>w/o MOUSE</sub> | 0.666 | <u>0.330</u> | 0.652 | 0.637 | 0.218 | <u>0.673</u> |
| SPROF-GO | <u>0.686</u> | **0.332** | **0.676** | **0.662** | **0.231** | **0.686** |

*Note:* SPROF-GO<sub>species</sub> denotes the model trained with proteins of the target species only, and SPROF-GO<sub>w/o species</sub> denotes the model trained with proteins of all species except the target species. Bold and underlined fonts indicate the best and second-best results, respectively.

## Reference

1. Eisenberg D, Marcotte EM, Xenarios I et al. Protein function in the post-genomic era, Nature 2000;405:823–826.

2. Costanzo M, VanderSluis B, Koch EN et al. A global genetic interaction network maps a wiring diagram of cellular function, Science 2016;353.

3. The UniProt Consortium. UniProt: the universal protein knowledgebase in 2021, Nucleic acids research 2021;49:D480–D489.

4. Cruz LM, Trefflich S, Weiss VA et al. Protein function prediction, Functional Genomics 2017:55–75.

5. Radivojac P, Clark WT, Oron TR et al. A large-scale evaluation of computational protein function prediction, Nature methods 2013;10:221–227.

6. Ashburner M, Ball CA, Blake JA et al. Gene ontology: tool for the unification of biology, Nature genetics 2000;25:25–29.

7. Obozinski G, Lanckriet G, Grant C et al. Consistent probabilistic outputs for protein function prediction, Genome Biology 2008;9:1–19.

8. Jiang Y, Oron TR, Clark WT et al. An expanded evaluation of protein function prediction methods shows an improvement in accuracy, Genome Biology 2016;17:1–19.

9. Zhou N, Jiang Y, Bergquist TR et al. The CAFA challenge reports improved protein function prediction and new functional annotations for hundreds of genes through experimental screens, Genome Biology 2019;20:1–23.

10. Conesa A, Götz S, García-Gómez JM et al. Blast2GO: a universal tool for annotation, visualization and analysis in functional genomics research, Bioinformatics 2005;21:3674–3676.

11. You R, Zhang Z, Xiong Y et al. GOLabeler: improving sequence-based large-scale protein function prediction by learning to rank, Bioinformatics 2018;34:2465–2473.

12. Altschul SF, Madden TL, Schäffer AA et al. Gapped BLAST and PSI-BLAST: a new generation of protein database search programs, Nucleic acids research 1997;25:3389–3402.

13. Jones P, Binns D, Chang H-Y et al. InterProScan 5: genome-scale protein function classification, Bioinformatics 2014;30:1236–1240.

14. Kulmanov M, Hoehndorf R. DeepGOPlus: improved protein function prediction from sequence, Bioinformatics 2020;36:422–429.

15. Cao Y, Shen Y. TALE: Transformer-based protein function Annotation with joint sequence–Label Embedding, Bioinformatics 2021;37:2825–2833.

16. Gligorijević V, Renfrew PD, Kosciolek T et al. Structure-based protein function prediction using graph convolutional networks, Nature communications 2021;12:1–14.

17. Lai B, Xu J. Accurate protein function prediction via graph attention networks with predicted structure information, Briefings in Bioinformatics 2022;23:bbab502.

18. Oliver S. Guilt-by-association goes global, Nature 2000;403:601–602.

19. You R, Yao S, Xiong Y et al. NetGO: improving large-scale protein function prediction with massive network information, Nucleic acids research 2019;47:W379–W387.

20. Szklarczyk D, Gable AL, Nastou KC et al. The STRING database in 2021: customizable protein–protein networks, and functional characterization of user-uploaded gene/measurement sets, Nucleic acids research 2021;49:D605–D612.

21. Kulmanov M, Khan MA, Hoehndorf R. DeepGO: predicting protein functions from sequence and interactions using a deep ontology-aware classifier, Bioinformatics 2018;34:660–668.

22. Torres M, Yang H, Romero AE et al. Protein function prediction for newly sequenced organisms, Nature Machine Intelligence 2021;3:1050–1060.

23. You R, Yao S, Mamitsuka H et al. DeepGraphGO: graph neural network for large-scale, multispecies protein function prediction, Bioinformatics 2021;37:i262–i271.

24. You R, Huang X, Zhu S. DeepText2GO: Improving large-scale protein function prediction with deep semantic text representation, Methods 2018;145:82–90.

25. Yao S, You R, Wang S et al. NetGO 2.0: improving large-scale protein function prediction with massive sequence, text, domain, family and network information, Nucleic acids research 2021;49:W469–W475.

26. Rives A, Meier J, Sercu T et al. Biological structure and function emerge from scaling unsupervised learning to 250 million protein sequences, Proceedings of the National Academy of Sciences 2021;118.

27. Elnaggar A, Heinzinger M, Dallago C et al. ProtTrans: Towards Cracking the Language of Lifes Code Through Self-Supervised Deep Learning and High Performance Computing, IEEE transactions on pattern analysis and machine intelligence 2021.

28. Unsal S, Atas H, Albayrak M et al. Learning functional properties of proteins with language models, Nature Machine Intelligence 2022;4:227–245.

29. Yuan Q, Chen S, Wang Y et al. Alignment-free metal ion-binding site prediction from protein sequence through pretrained language model and multi-task learning, Briefings in Bioinformatics 2022.

30. Cowen L, Ideker T, Raphael BJ et al. Network propagation: a universal amplifier of genetic associations, Nature Reviews Genetics 2017;18:551–562.

31. Boutet E, Lieberherr D, Tognolli M et al. UniProtKB/Swiss-Prot, the manually annotated section of the UniProt KnowledgeBase: how to use the entry view. Plant Bioinformatics. Springer, 2016, 23–54.

32. Huntley RP, Sawford T, Mutowo-Meullenet P et al. The GOA database: gene ontology annotation updates for 2015, Nucleic acids research 2015;43:D1057–D1063.

33. Raffel C, Shazeer N, Roberts A et al. Exploring the Limits of Transfer Learning with a Unified Text-to-Text Transformer, Journal of machine learning research 2020;21:1–67.

34. Steinegger M, Mirdita M, Söding J. Protein-level assembly increases protein sequence recovery from metagenomic samples manyfold, Nature methods 2019;16:603–606.

35. Suzek BE, Huang H, McGarvey P et al. UniRef: comprehensive and non-redundant UniProt reference clusters, Bioinformatics 2007;23:1282–1288.

36. Kenton JDM-WC, Toutanova LK. BERT: Pre-training of Deep Bidirectional Transformers for Language Understanding. In: Proceedings of NAACL-HLT. 2019, p. 4171–4186.

37. Ba JL, Kiros JR, Hinton GE. Layer Normalization, stat 2016;1050:21.

38. Srivastava N, Hinton G, Krizhevsky A et al. Dropout: a simple way to prevent neural networks from overfitting, The journal of machine learning research 2014;15:1929–1958.

39. Giunchiglia E, Lukasiewicz T. Coherent Hierarchical Multi-Label Classification Networks. In: Advances in neural information processing systems. 2020, p. 9662–9673. Curran Associates, Inc.

40. Buchfink B, Xie C, Huson DH. Fast and sensitive protein alignment using DIAMOND, Nature methods 2015;12:59–60.

41. Kingma DP, Ba J. Adam: A Method for Stochastic Optimization. In: 3rd International Conference on Learning Representations (Poster). 2015.

42. Paszke A, Gross S, Massa F et al. Pytorch: An imperative style, high-performance deep learning library, Advances in neural information processing systems 2019;32:8026–8037.

43. Davis J, Goadrich M. The relationship between Precision-Recall and ROC curves. In: Proceedings of the 23rd international conference on Machine learning. 2006, p. 233–240.

44. Saito T, Rehmsmeier M. The precision-recall plot is more informative than the ROC plot when evaluating binary classifiers on imbalanced datasets, PloS one 2015;10:e0118432.

45. Fukata M, Fukata Y, Adesnik H et al. Identification of PSD-95 palmitoylating enzymes, Neuron 2004;44:987–996.

46. Yamamoto M, Ko LJ, Leonard MW et al. Activity and tissue-specific expression of the transcription factor NF-E1 multigene family, Genes & development 1990;4:1650–1662.

47. Yuan Q, Chen J, Zhao H et al. Structure-aware protein–protein interaction site prediction using deep graph convolutional network, Bioinformatics 2021;38:125–132.

48. Yuan Q, Chen S, Rao J et al. AlphaFold2-aware protein–DNA binding site prediction using graph transformer, Briefings in Bioinformatics 2022;23:bbab564.

49. Varadi M, Anyango S, Deshpande M et al. AlphaFold Protein Structure Database: massively expanding the structural coverage of protein-sequence space with high-accuracy models, Nucleic acids research 2022;50:D439–D444.

50. Lin Z, Akin H, Rao R et al. Evolutionary-scale prediction of atomic level protein structure with a language model, bioRxiv 2022.

51. Chen T, Kornblith S, Norouzi M et al. A simple framework for contrastive learning of visual representations. In: International conference on machine learning. 2020, p. 1597–1607. PMLR.

52. Zheng S, Rao J, Song Y et al. PharmKG: a dedicated knowledge graph benchmark for bomedical data mining, Briefings in Bioinformatics 2021;22:bbaa344.

## Reference

1. Elnaggar A, Heinzinger M, Dallago C et al. ProtTrans: Towards Cracking the Language of Lifes Code Through Self-Supervised Deep Learning and High Performance Computing, IEEE transactions on pattern analysis and machine intelligence 2021.

2. Raffel C, Shazeer N, Roberts A et al. Exploring the Limits of Transfer Learning with a Unified Text-to-Text Transformer, Journal of machine learning research 2020;21:1–67.

3. Steinegger M, Mirdita M, Söding J. Protein-level assembly increases protein sequence recovery from metagenomic samples manyfold, Nature methods 2019;16:603–606.

4. Suzek BE, Huang H, McGarvey P et al. UniRef: comprehensive and non-redundant UniProt reference clusters, Bioinformatics 2007;23:1282–1288.

5. Kenton JDM-WC, Toutanova LK. BERT: Pre-training of Deep Bidirectional Transformers for Language Understanding. In: Proceedings of NAACL-HLT. 2019, p. 4171–4186.

6. Zhou N, Jiang Y, Bergquist TR et al. The CAFA challenge reports improved protein function prediction and new functional annotations for hundreds of genes through experimental screens, Genome Biology 2019;20:1–23.

7. Davis J, Goadrich M. The relationship between Precision-Recall and ROC curves. In: Proceedings of the 23rd international conference on Machine learning. 2006, p. 233–240.

8. Saito T, Rehmsmeier M. The precision-recall plot is more informative than the ROC plot when evaluating binary classifiers on imbalanced datasets, PloS one 2015;10:e0118432.

9. Kulmanov M, Hoehndorf R. DeepGOPlus: improved protein function prediction from sequence, Bioinformatics 2020;36:422–429.

10. You R, Yao S, Mamitsuka H et al. DeepGraphGO: graph neural network for large-scale, multispecies protein function prediction, Bioinformatics 2021;37:i262–i271.

11. Altschul SF, Madden TL, Schäffer AA et al. Gapped BLAST and PSI-BLAST: a new generation of protein database search programs, Nucleic acids research 1997;25:3389–3402.

12. Mitchell AL, Attwood TK, Babbitt PC et al. InterPro in 2019: improving coverage, classification and access to protein sequence annotations, Nucleic acids research 2019;47:D351–D360.

13. Mistry J, Chuguransky S, Williams L et al. Pfam: The protein families database in 2021, Nucleic acids research 2021;49:D412–D419.

14. Dawson NL, Lewis TE, Das S et al. CATH: an expanded resource to predict protein function through structure and sequence, Nucleic acids research 2017;45:D289–D295.

15. Lu S, Wang J, Chitsaz F et al. CDD/SPARCLE: the conserved domain database in 2020, Nucleic acids research 2020;48:D265–D268.

16. Jones P, Binns D, Chang H-Y et al. InterProScan 5: genome-scale protein function classification, Bioinformatics 2014;30:1236–1240.

17. Buchfink B, Xie C, Huson DH. Fast and sensitive protein alignment using DIAMOND, Nature methods 2015;12:59–60.

18. Szklarczyk D, Gable AL, Nastou KC et al. The STRING database in 2021: customizable protein–protein networks, and functional characterization of user-uploaded gene/measurement sets, Nucleic acids research 2021;49:D605–D612.

